# The contribution of small protected areas to the big picture of forest conservation and restoration in Sweden

**DOI:** 10.1101/2024.05.27.596035

**Authors:** Johan Svensson, Andres Lopez-Peinado, Bengt Gunnar Jonsson, Navinder J Singh

## Abstract

In forest regions worldwide, industrial forestry has left fragments of natural forests behind. This challenges biodiversity conservation and calls for ecological restoration for sustainable forest management and conservation. The functionality of protected areas need to be improved and forest ecosystems set in a state that better favors biodiversity, resilience and provisioning of ecosystem services. Sweden contributes a substantial share of the European forests, with dominance of non-industrial forest ownership and extensive forestry footprint, and hence with immediate need for advanced conservation and restoration. Protection through voluntary nature conservation agreements and regulated biotope protection areas exists since the 1990s, with schemes involving economic compensation to landowners to facilitate conservation and restoration. Across entire Sweden and all ecoregions, we assessed their accumulated capacity over a 30-year period, including forest types protected, type of restoration management, rotation intervals, and selection of tree species. These nearly 14,000 different areas covering over 70,000ha are small, ranging in size from 5ha but rarely larger than 20ha. Their contribution is important, particularly in south Sweden with low and fragmented forest cover among many different owners. Active restoration dominates over passive set asides, coniferous forest types are less represented than more rare forest types, many different tree species are favored, and different restoration types occur but with few types dominating. In recognizing their critical importance, we find that the practices are narrow and repetitive, and that a greater restoration diversification is needed. The decreasing trend in protection is alarming since these contribute key forest type representativeness and functionality.

## Introduction

The contribution of small-size protected areas in securing functional forest ecosystems and preserving biodiversity is generally considered as more limited in comparison with large-size areas (e.g., Fletcher et al. 2018; Timmers et al. 2022), as originally based on the theory of island biogeography (Mac Arthur & Wilson 1967). In many forest regions worldwide, however, scattered land ownership and extensive industrial forestry has resulted in fragmentation and loss of natural forest habitats including primary and old growth forest, leaving only small patches of high conservation value forests behind (Heino et al. 2015; Svensson et al. 2019; Sabatini et al. 2020). Under these circumstances, conservation planning necessarily relies on small-size protected areas, with their contribution to biodiversity conservation being recognized as valuable (Wintle et al. 2019; Fahrig 2020; Riva et al. 2022). To be ecologically functional, they need to collectively provide a functional network of forests that together with larger protected areas ensure landscape connectivity, biodiversity and natural pools of ecosystem services (Orlikowska et al. 2020; Strassburg et al. 2020).

In regions and countries with extensive and long-term industrial forestry impact and fragmentation, ecological restoration is required to meet agreed international conservation goals; e.g. the Kunming-Montreal Global Biodiversity Framework (CBD 2022) and the EU Biodiversity Strategy (EC 2020). Thereby, restoration becomes a key component in forest and forest landscape planning approaches (Chazdon et al. 2016; Besseau et al. 2018; Guiterrez et al. 2022). Where private and scattered forest ownership dominates, such as in many European countries, coherent and effective conservation planning is additionally challenging (Tiebel et al. 2022). Furthermore, these challenges are multi-facetted since diversification and multiple aspects of forest land use (Knoke et al. 2017; Felton et al. 2020; Blattert et al. 2023) point at restoration needs beyond only improving nature conservation attributes (Vallecillo et al. 2018; Svensson et al. 2023a). Hence, well-informed implementation of the EU Nature Restoration Law (EC 2024; Bou Dagher Kharrat et al. 2023) and other high-level initiatives and agreements, e.g., the UN Decade on Ecosystem Restoration (UN 2019) and the CBD Ecosystem Restoration target on restoring 30% of degraded ecosystems to 2030 (CBD 2022), is critically needed.

The north European forest landscape has undergone extensive transformation during the 20^th^ century (e.g., Pohjanmies et al. 2017). Vast areas are converted into plantation-like forests resulting in severe habitat fragmentation and loss of natural and socio-cultural capital (IPBES 2018; Fleishman et al. 2022; Sullivan et al. 2023; Tadesco et al. 2023). With 28 million hectare forestlands (SLU 2023), Sweden contributes about 12% (Forest Europe 2020) of European forests. Despite an increased policy attention on environmental consideration since the 1990s (Beland-Lindahl et al. 2017), the outcome of a strongly dominating biomass-yield forestry model (e.g., Jonsson et al. 2019) is that Sweden does not meet agreed national, European, and international forest conservation and biodiversity goals (Angelstam et al. 2020). Restoration is, hence, necessary as a component in the national forestry model in order to secure transformation of large areas of forests, improve ecological functionality and favorable conservation status, diversify forest conditions, and improve biodiversity and resilience (Halme et al. 2013).

Furthermore, restoration is required at multiple scales, i.e. from stands to landscapes, across all ecoregions and native forest habitat types (Svensson et al. 2023a). Since protected forest areas are strongly biased to the hinterland mountainous region in Sweden, 58% in the mountain region compared with 4% of all forestland elsewhere (Statistics Sweden 2023), representing a narrow segment of habitat types, further protection needs to expand outside this region to conserve a representative share of all remaining habitat types (Cf. Riva et al. 2022). The national share of forestland outside the mountain region is large, 91% and 95% of all forestland and forest of interest for commercial forestry, respectively (Statistics Sweden 2023), as is the share of non-industrial private forest (NIPF) owners. In Sweden, 48% of the forestland of interest for active forestry is on NIPF land ownership (Statistics Sweden 2019). With the marked industrial forestry footprint on forests outside the mountain region (Bubnicki et al. 2024), further conservation as well as restoration will need to involve NIPF land-ownership.

Nature Conservation Agreements (NCA) and Biotope Protection Areas (BPA) are recognized as formal forest protection instruments in Sweden, with NCA being voluntary and BPA being regulated by the State (Statistics Sweden 2023); i.e. can be forced by State authorities. Both instruments are applied on NIPF land ownership, and both cover small-size areas. According to official statistics 1994 up to 2023 (Swedish Forest Agency 2024), these together include 14,432 different patches and in total 74,773 hectare, with an average area of 7.2 (NCA) and 3.9 hectare (BPA), respectively. Both NCA and BPA are based on an identification of a woodland key habitat, where the structure, history, physical conditions and presence of species indicate high forest conservation values (Timonen et al. 2010). Furthermore, these small and scattered NCA and BPA protection patches include representative forest habitat types from the nemoral biome in the south to the boreal and subalpine biomes in the north, contributing to the national, European and global conservation agenda. Whereas a share of the area is set aside for natural free development, another share is allocated for restoration and active conservation management. Hence, these instruments represent a national-scale ecological restoration approach with specified management activities to enhance the conservation value (e.g. Riva et al. 2022), improve ecological functionality (e.g., Brennan et al. 2022), contribute to a broader array of ecosystem services (e.g., Peura et al. 2018), and restore biotic and abiotic interactions in forest ecosystems.

In this study we assessed the contribution to forest protection and restoration of NCAs and BPAs in Sweden. With basic data on location, surface and forest area protected over a 30-year time period, we focused our analyses on the ecological aspects of protection and restoration; i.e. which forest types are protected, what type of ecological restoration is applied, at which rotation intervals, and which tree species are favored vs. disfavored. While NCA and BPA instruments have existed for a long time, their extent, dynamics, values, and contribution to comprehensive landscape-scale conservation, has not been explored. The scientific community, policymakers, landowners and managers lack a comprehensive understanding of their role and potential as a formal protection instrument. We discuss the outcome and potential of small-size protection and restoration across a broad biophysical gradient, to what extent these contribute to fulfilling national and international environmental goals, and the opportunities that arises for diversifying forestry practices and facilitating more resilient forest ecosystem. Finally, we also discuss the socio-economic aspects of NCA as a voluntary and more general protection instrument vs. BPA as a strict biodiversity protection instrument, and the implications of our analyses in the context of scattered and small-size conservation and restoration by private actors.

## Methods

### Study area description

Sweden is dominated by forestlands, i.e. 69% (Statistics Sweden 2019) of the land base, from the nemoral biome and habitat types in the south to boreal and subalpine in the north, and with pronounced biogeographic and socio-economic gradients. Scots pine (*Pinus sylvestris*) forests dominate (40%), followed by Norway spruce (*Picea abies*) forests 28%, mixed coniferous forests 13% and mixed coniferous and deciduous forests 7% (Angelstam & Manton 2021). Deciduous forests cover 7%, including 1.1 million ha of subalpine mountain birch (*Betula pubescens* ssp. *czerepanovii*) alpine tree line forests of the Scandinavian Mountain Range (Hedenås et al. 2016). Hardwood deciduous forests, mainly with oak (mainly *Quercus robur*) and beech (*Fagus sylvatica*) occur frequently in southern nemoral and boreonemoral regions Sweden.

Non-industrial private forest owners (NIPF) are the dominant landowners with 48% of the forest area of interest for the commercial forestry; i.e. forest on sites that on average over a rotation period annually can produce a wood biomass growth ≥ 1m^3^/ha (SLU 2023). Private forest companies owns 24%, the state and state controlled companies 20%, and other private and public owners 8% of the forestland (Statistics Sweden 2019). Commonly, NIPF-ownership occurs in particular in south, central and along the northeast coast where urban and settlement areas are abundant, and at lower terrain with more fertile mineral soils (Statistics Sweden 2019; SLU 2023). Hence, NIPF-ownership is generally located closer to forest industries and on land with higher site productivity and forest production capacity. Consequently, NIPF-owners provide 61% of the raw material to the Swedish forest industry (SLU 2023), and are thus critically important for the Swedish forest industry. In total 311,000 different NIPF-owners (physical persons) were registered in 2022 (Swedish Forest Agency, 2024). This ownership is within 232,000 management units with an average and median forest area of 34 ha and 12 ha, respectively (Swedish Forest Agency 2021). Hence, the majority of the management units are small by comparison; 32% ≤5ha, 63% ≤5ha, and 83% ≤50ha (Swedish Forest Agency 2024).

As of December 31, 2022, totaling 2.4 million ha (9%) forestland is formally protected (Statistics Sweden 2023). However, the protected area is highly biased geographically and with respect to targets of representative protection of naturally occurring forest habitats, cf. the Montreal-Kunming CBD Framework (CBD 2022); 58% of the forestland is protected in the mountain region, compared with 5%, 3%, 4% and 4%, respectively, in the north boreal, south boreal, boreonemoral and nemoral regions protected (Statistics Sweden 2023). Consequently, forest protection is highly concentrated to a segment of the native habitat types and to habitat in subalpine and generally less rich and productive site types.

We have analyzed NCA and BPA distribution across Sweden with stratification into four ecoregions, following Statistics Sweden (2023) but with the mountain and mountain foothills region included in the north boreal and south boreal regions (Fig. 1). This is owing to the limited forest areas and abundance of NCA and BPA in the mountain and mountain foothills region, and to our focus on restoration management.

**Figure 1.**
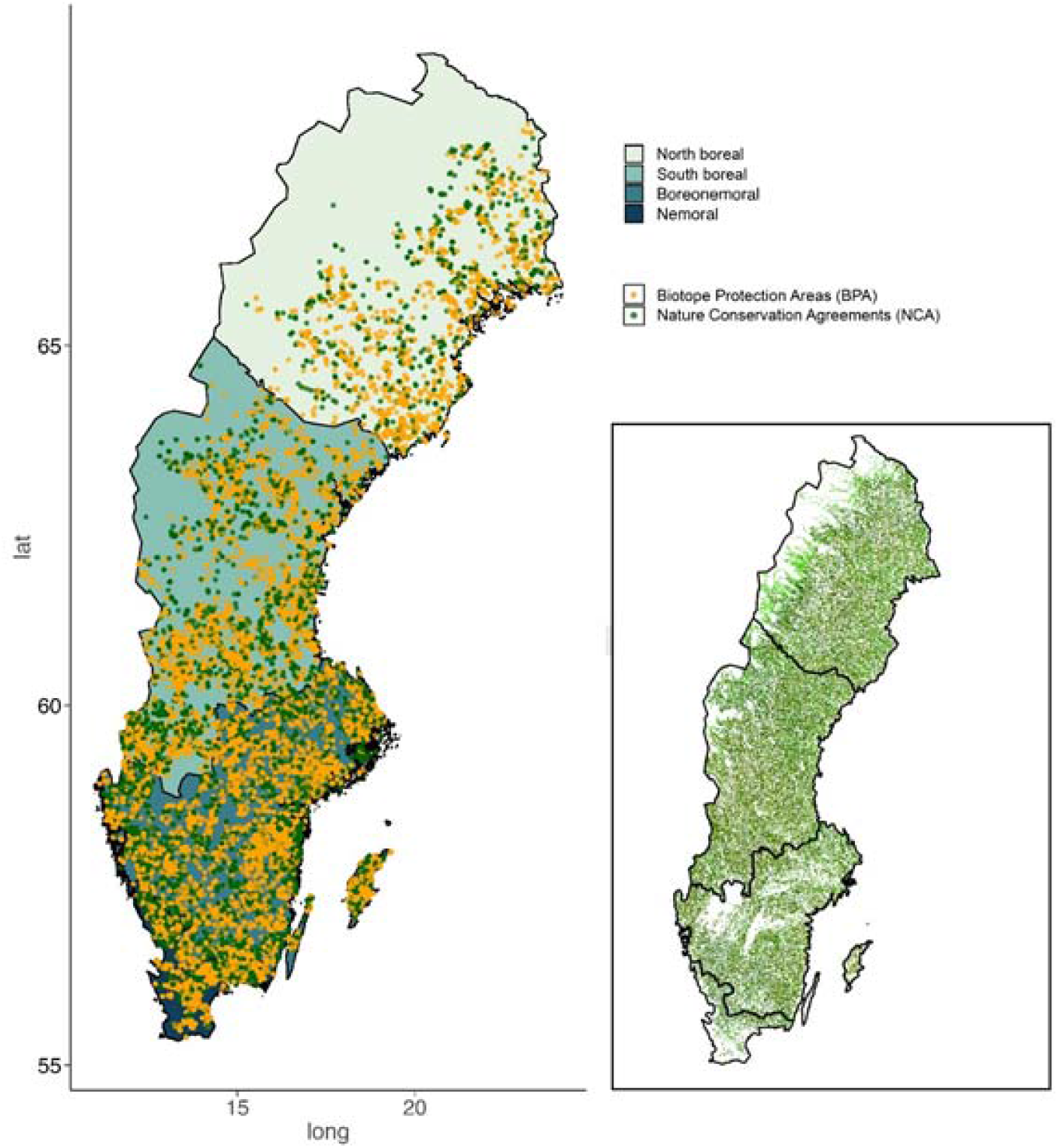
Spatial distribution of all Nature Conservation Agreements (NCA; from the first established October 1, 1993) in green, and all Biotope Protection Areas (BPA; from the first established February 28, 1994), in yellow, up to 2023 (June 6). Displayed on the four ecoregions applied in this study; from north to south and darker color: North boreal, South boreal, Boreonemoral, Nemoral. The insert map shows the forest cover in Sweden (Swedish Environmental Protection Agency 2023).

### Nature conservation Agreements and Biotope Protection Areas

Nature conservation agreements are voluntary agreements on forestland according to the Swedish Land Code (1970) between a landowner and the State, represented by the Swedish Forest Agency (Statistics Sweden 2018). The agreement states that the landowner, within a certain time-period and with a certain level of economic compensation, take consideration to nature conservation values on top of the legal requirement by the Swedish Forestry Act (Swedish Forest Agency 2023). The maximum time-period of the agreement is 50 years, but agreements with shorter time-periods can be made. The standard economic compensation is 60% of the merchantable value of net standing wood biomass (i.e. reduced by projected harvesting and logging costs) for a 50-year agreement, but less for shorter agreements (Statistics Sweden 2018). The NCA formal protection instrument has been applied also on larger forest landscapes with State and private forest companies, e.g. with Sveaskog state forest company on 37 ecoparks covering between 1,100 and 22,000 ha (Bergman & Gustafsson 2020). These landscape NCAs are not included in this study.

Biotope protection areas are, like nature reserves, stronger regulations in the sense that the State can enforce the protection following the Environmental Code (1998), either by expropriating the land or by establishing formal decision rights for management of the above ground vegetation. Normally, areas larger than around 20 ha become nature reserves, whereas smaller areas become BPAs. The compensation to land owner is regulated to a one-time payment of a sum equal to 125% of the reduced market value for the management unit in question (Statistics Sweden 2018).

As state-controlled formal protection instruments, NCA represents a soft regulation and BPA a strict regulation, respectively. Both protection instruments contribute with representativeness and diversity of forest habitat types to the national conservation network on non-industrial private forest (NIPF) ownership. Commonly. NIPF-ownership dominate closer urban and settlement areas, generally with higher site productivity and thus potentially higher biodiversity characteristics; the site wood biomass production capacity on NIPF ownership averages 6.3 compared with 5.5 m^3^/ha,year as a national average (SLU 2023). Furthermore, NIPF-ownership generally also include forests with a longer and more intense land-use history with open and semi-open conditions, livestock grazing and small-scale forestry (Statistics Sweden 2019). Thus, the present forest conservation values may be more diverse and multi-facetted, calling for flexible conservation strategies as well as for diversified active restoration activities.

### Data and analyses

The data consist of all NCA from the first established in 1993 (October 1) and all BPA from the first established in 1994 (February 28) up to 2023 (June 26), in a database provided by the Swedish Forest Agency that contains in total 13,779 objects covering 71,164hectare (Table 1). A very large share of the total protected area consists of productive forest of potential interest for active forestry; 97% in BPAs and 90% in NCAs.

**Table 1.** Forest area (Ha), number of objects (No) and total area (Ha) of Nature Conservation Agreements (NCA) and Biotope Protection Areas (BPA), presented per ecoregion and in total.

|  | Forest area* | Protected forest area* | NCA |  | BPA |  |
| --- | --- | --- | --- | --- | --- | --- |
|  | Ha | Ha | No | Ha | No | Ha |
| North boreal | 8,740,400 | 1,564,300 | 591 | 5,685 | 1,042 | 4,591 |
| South boreal | 10,300,500 | 514,200 | 1,847 | 13,469 | 2,756 | 11,314 |
| Boreonemoral | 6,909,100 | 277,000 | 2,384 | 15,766 | 3,811 | 14,218 |
| Nemoral | 957,900 | 42,800 | 457 | 3,133 | 891 | 2,988 |
| Total | 26,907,900 | 2,398,300 | 5,279 | 38,053 | 8,500 | 33,111 |

\* Data from Statistics Sweden 2023. Statistics Sweden (2023) report 38,000 ha NCA and 31,900 ha BPA on forestland up to December 2022, i.e. total area provided here include area up to June 2023 included a fraction of other land cover than forests.

Applied data for this study include for all objects: Polygon with centroid position (long., lat.), total area, forest area of interest for production forestry (see above), agreement date, area of restoration management activity, type of restoration management activity, target tree species, and return interval of restoration management activity. Type of restoration management activity in the database includes in total 30 different types, whereof 13 combined types were constructed: 1) Forest floor and hydrology; 2) Shrub layer; 3) Tree regeneration; 4) Tree age heterogeneity; 5) Removal of tree species; 6) Thinning from above; 7) Gap cutting; 8) Dead wood; 9) Prescribed burning; 10) Forest edge; 11) Cultural and recreational values; 12) Other; 13) Set aside (for free development). Restoration type combinations are outlined in the supplementary material (Table S1). For the analysis on return interval of restoration management activity, only types 3, 5, 6, 7, 8 and 9 were used, to reflect management activities within forest stands. Type 4, tree age heterogeneity, was excluded in this analysis since the data compilation showed highly varied restoration goals both within the forest area and for other land covers and structures within the total area, and since a substantial share of the restoration concerned removing forest to re-create pasture conditions, blocking ditches, etc. The analyses on favored and dis-favored tree-species concerned only categories 3, 5 and 6, where restoration activities are directed specifically to selected tree species. For the analysis on favored tree species we applied subtypes within type 3 (Tree specific regeneration management) and type 6 (Favor selected tree species and tree groups in canopy layer), where restoration area can be defined to specific species. Area calculations for restoration management activities overlap if different activities are planned on the same area. Thus, these areas are absolute for any activity but can not be totaled across several activities. Thereby, comparison with area set aside for natural, free development, where such overlap do not exist, can be made.

As complementary wall-to-wall data on forest type, i.e. dominating tree species, we used the publicly available National Land Cover data (NLC; Swedish Environmental Protection Agency 2023) forest types on 10x10m resolution, with in total eight types on either wet, organic or dry, mineral parent material. In this study, we applied the following categories: 1) Pine forest; 2) Spruce forest; 3) Mixed coniferous forest; 4) Mixed forest; 5) Deciduous forest; 6) Deciduous hardwood forest. The latter category combined NLC-categories “hardwood deciduous forest” and “hardwood deciduous forest with trivial deciduous forest”. Pine, spruce and deciduous forests are defined as forest with ≥70% of the tree crown cover by that or those species. The NLC-category “temporarily not forest”, was not applied.

All data compilation and analyses were made in R (R Core Team 2023) and QGIS (QGIS.org 2024).

## Results

In total since 1993, 5,279 NCAs have been established on 38,053ha, and in total since 1994, 8,500 BPAs on 3,311 ha (Fig. 2a). With a sharp increase in the end of the 1990s and beginning of 2000s, the largest share of the BPAs were protected between 2000 and 2007, and of the NCAs between 2002 and 2009. The number of objects and areas are decreasing since 2006, whereas the average size of both instruments are increasing significantly over time (Fig. S2). NCAs are generally larger than BPAs (Fig. 2b). Their size-distribution shows a concentration to areas up to ca. 5 ha, with NCA and BPA size-ranges generally up to ca. 20ha and 7ha, respectively, and with larger objects being infrequent. The largest NCA in the dataset is 221ha and the largest BPA is 25ha.

**Figure 2.**
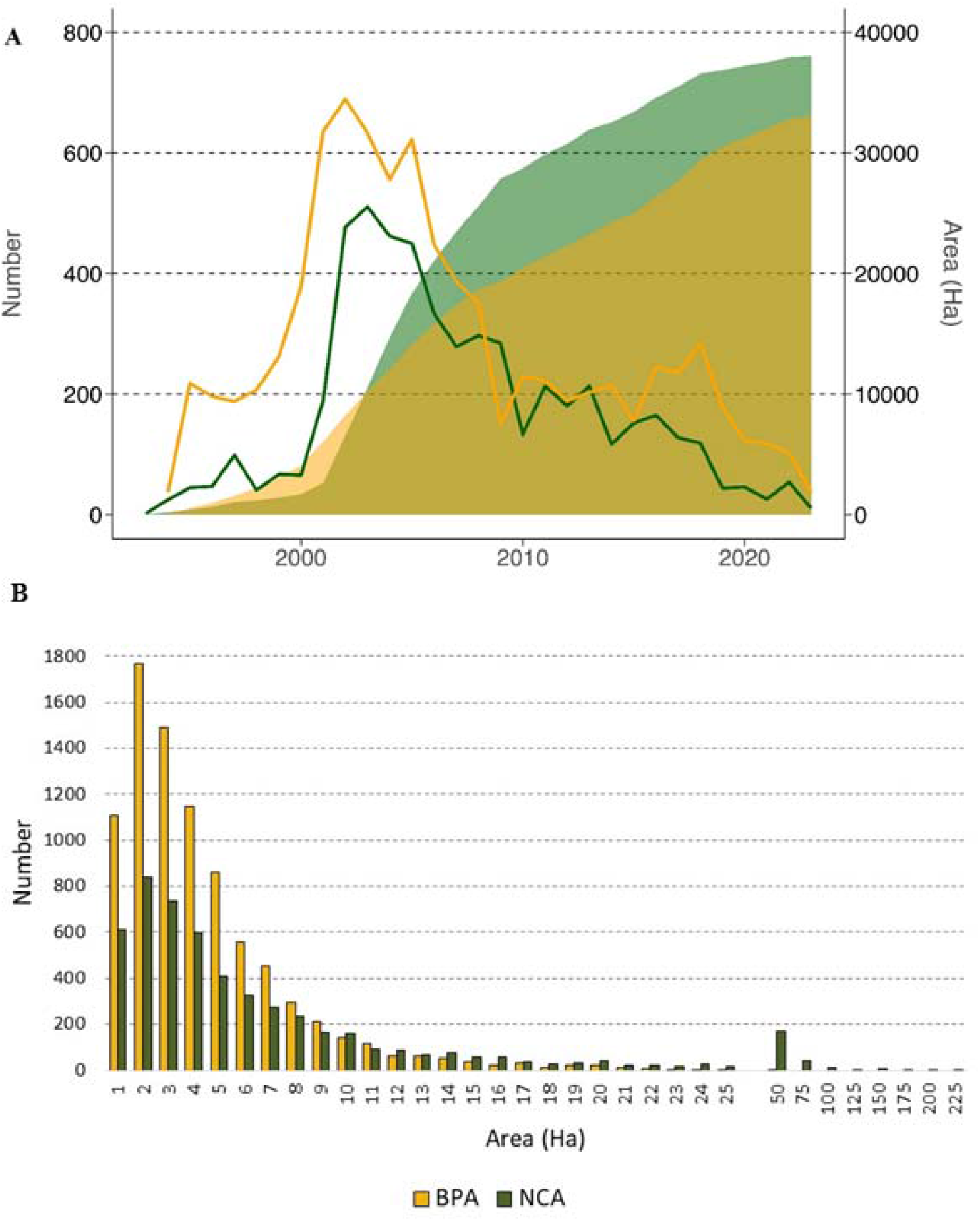
Number per year and accumulated area of Nature Conservation Agreements (NCA) and Biotope Protection Areas (BPA) over time (A), and the size distribution of NCA and BPA (B). The size distribution is per 1ha classes (with highest class-value shown on the axis) up to 24.1–25.0ha, followed by 25.1–50.0ha, 50.1–75.0ha, etc. Mean, median and standard deviation statistics for NCA is 7.2ha, 3.8ha and 12.3ha, and for BPA 3.9ha, 3.0ha and 3.4ha.

Compared with the total forest area per ecoregion, NCA and BPA cover small proportions; from 0.07% (NCA) and 0,36% (BPA) in the north boreal ecoregion, to 0.33% (NCA) and 0.31% (BPA) in the nemoral (see Table 1). Compared with the total protected forest area, however, the two instruments combined cover 0.7%, 4.8%, 10.8% and 14.3% in the north boreal, south boreal, boreonemoral and nemoral ecoregions, respectively.

Protection of deciduous hardwood forests dominate in the nemoral ecoregion, deciduous forests and pine forest in the boreonemoral ecoregion, and pine and spruce forests in the south and north boreal ecoregions (Fig. 3, Table S3), reflecting the dominant forest types in these ecoregions. Forest on wet, organic soils occur at low abundance generally. Overall across Sweden, pine (in total 19,964ha) and spruce forests (18,830ha) are dominating in NCAs and BPAs, followed by deciduous forests (11,888ha), mixed forests (9,522ha), mixed coniferous forests (8,060ha), and deciduous hardwood forest (7,447ha).

**Figure 3.**
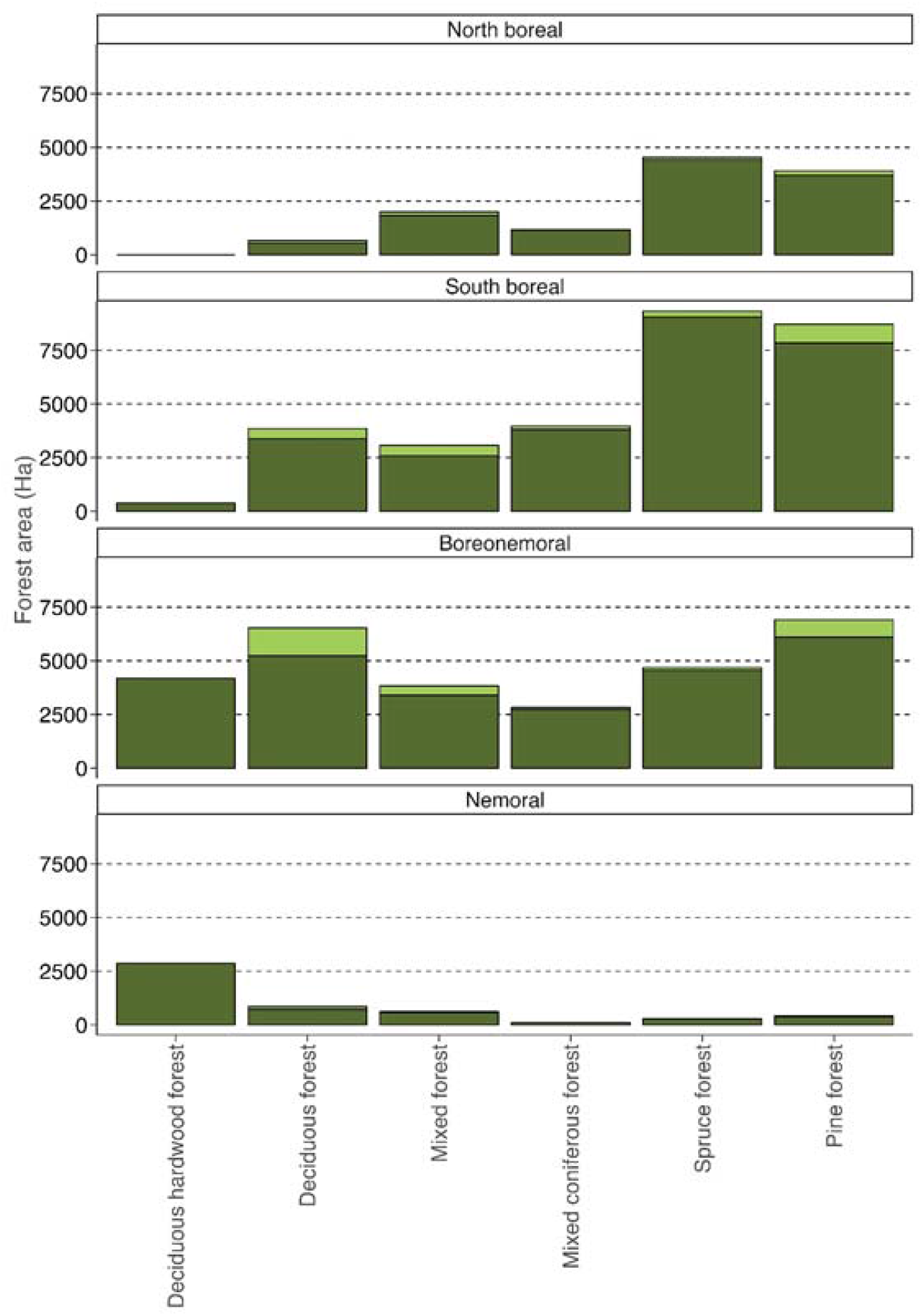
Area (ha) of forest types in Nature Conservation Agreements and Biotope Protection Areas (BPA) combined, presented per ecoregion. Surfaces on wetland are in light green color and not on wetland in dark green color. Land cover data extracted from Swedish Protection Agency (2023).

The largest area is set aside for natural, free development, in total 30,585ha, and in particular in the south and north boreal ecoregion (Fig. 4, Table S4). For restoration management activities within forest stands, removal of tree species and thinning from above dominate in the nemoral and boreonemoral ecoregions, and together with prescribed burning in the south boreal ecoregion. Prescribed burning dominated in the north boreal ecoregion. In total in Sweden, active restoration management concerning removal of tree species is performed on 9,420ha, and concerning thinning from above on 8,529ha. With exception of restoration of cultural and recreation values in particular in the boreonemoral region, other restoration activities than those oriented directly to trees, are infrequent. The combined area with active restoration management is 40,579ha, which equals 57% of the total NCA and BPA area.

**Figure 4.**
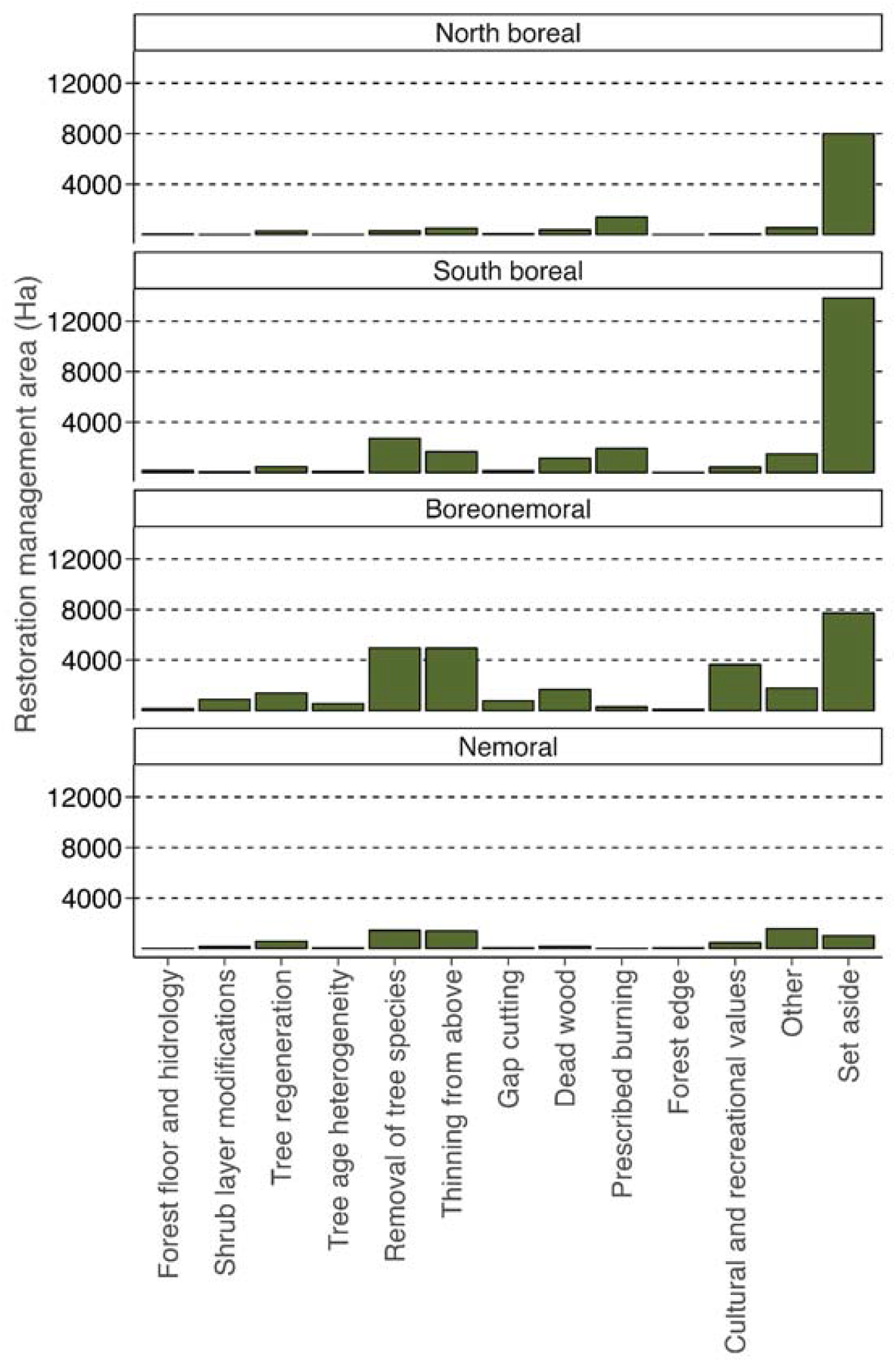
Area (ha) of restoration management types in Nature Conservation Agreements and Biotope Protection Areas (BPA) combined, presented per ecoregion.

For restoration management activities with forest stands, the planned re-occurring time interval for the activity is predominantly between 1 and 5 years, for all activities except prescribed burning where burning is planned at close to equal shares across all intervals (Table 2). For other activities, the longest interval of 30 to 50 years is infrequent.

**Table 2.** Proportion (%) of time interval (classes) between re-occurring restoration management activities.

|  | Time interval (year) |  |  |  |
| --- | --- | --- | --- | --- |
|  | 1-5 | 5-10 | 10-30 | 30-50 |
| Tree regeneration | 47 | 35 | 15 | 1 |
| Removal of tree species | 49 | 29 | 19 | 2 |
| Thinning from above | 50 | 30 | 17 | 1 |
| Gap cutting | 47 | 31 | 17 | 2 |
| Dead wood | 50 | 34 | 14 | 1 |
| Prescribed burning | 27 | 28 | 19 | 24 |

Footnote: The calculations were based on forest area with different restoration management activities. A fraction (ca. 8% in total and 1-2% per activity) of the area contains no data on time interval. These areas are considered in the totals of each activity, thus, the table do not sum to 100% horizontally.

Finally, for restoration management activities with specific tree species selection, our results show that restoration favors diversity in tree species composition (Figure 5, Table S5). In total 24 different species are specifically selected to be favored in management of understory and regeneration as well as in management of the canopy layer. Few species dominate, however, in the canopy layer particularly aspen (*Populus tremula*), birch (*Betula sp*.), oak (*Quercus sp*.) and pine, and in the understory aspen, birch, oak, pine, rowan (*Sorbus aucuparia*) and willow (*Salix sp*.). Of these species, only oak is restricted to the nemoral and boreonemoral ecoregions, whereas the other species occur throughout the forest landscapes in Sweden. In comparison, also disfavored species can be identified, showing that 95% of the area with restoration type removal of tree species concern Norway spruce; harvesting of trees >15 cm (dbh; diameter at 1.3m) and cleaning in understory regeneration (Table S6).

**Figure 5.**
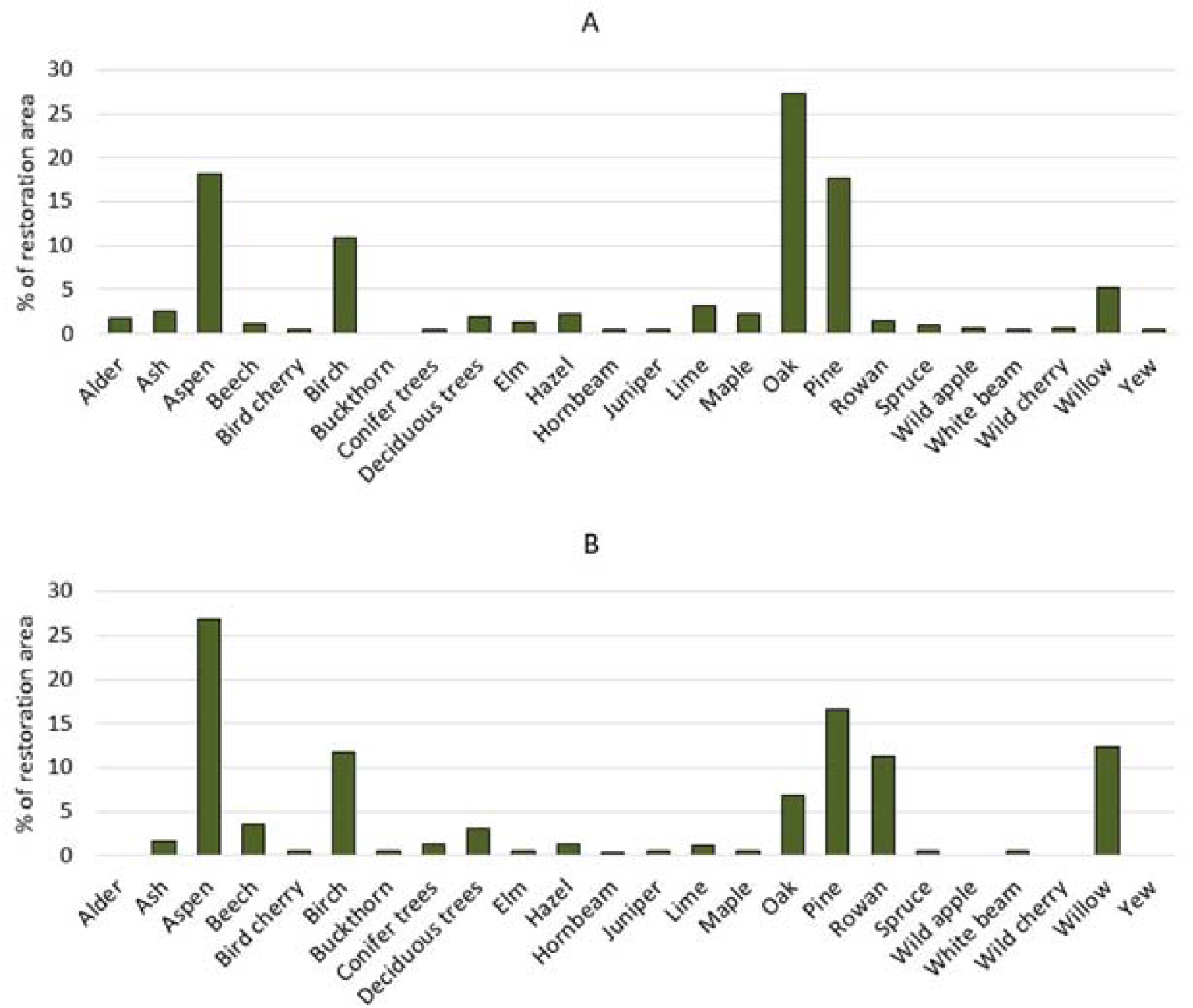
Favored tree and tall woody shrub species in canopy (A) and understory and regeneration (B), as percent of total restoration area of all favored tree and woody tall shrub species. For a), data was calculated in subtype Favor selected tree species and tree groups in canopy layer within type Thinning from above, and for b), data was calculated for subtype Tree specific regeneration management within type Tree regeneration (see Table S1). To be visible in the diagram, all occurring species was set to a minimum value of 0.5%, thus no bar show no occurrence. Calculations were made based on the area registered for each species by the total area registered for all species. Area without any species registered was not included. See Table S5 for data on area and Latin names.

## Discussion

The many small-size protected NCA- and BPA-areas complement larger protected areas such as nature reserves in the Swedish national, European Union and global conservation agenda, and contribute a diversity of forests in the larger landscape contexts. Despite that their overall area coverage is low relative to the vast forest cover in Sweden, their contribution to protected forest is substantial particularly in southern Sweden where forest cover proportionally less area, is generally more fragmented, and distributed among a large number of different ownership properties with a high share of non-industrial private owners (Statistics Sweden 2019; SLU 2023). As formal protection instruments, the growth of NCAs and BPAs since 1993 and 1994 up to 2023, showed a substantial increase in numbers of objects and area up to 2006, but following that, gradually fewer object and less area was established. The contribution in the most recent period, from 2020 to the end of June 2023, is as low as 137 NCAs contributing 1,178ha and 381 BPAs contributing 2,580ha. Since a higher deforestation rate globally has been detected among small forests compared with large and since a large number of small forests have been found to be disproportionally valuable for biodiversity conservation compared with large forests totaling same area (Riva et al. 2022), this decreasing trend is alarming. Further research is needed to define critical drivers, such as government funding, political initiatives, and willingness among landowners, that control this trend.

Our study focus on small-size protected areas, i.e. commonly up to ca. 5ha but with infrequent BPAs up to 25ha and NCAs up to more than 200ha. In comparison with, e.g., Brennan et al. (2022), where small protected areas was defined as those under 3,500 hectare, this is very small areas indeed. To draw parallels with countries with extensive networks of protected areas, such as, e.g., Costa Rica and Bolivia, the NCAs and BPAs are examples of private or community reserves often owned and managed by local communities, private landowners, or non-governmental organizations, where they play a crucial conservation role (Ferraro et al. 2013). Similar understandings about the substantial conservation capacity in private forest ownership can be made for Europe, e.g. for Germany (Tiebel et al. 2022). In Sweden, BPAs are similar to nature reserves in terms of a more strict biodiversity protection focus, whereby larger areas and areas with several different owners (i.e. 20ha or larger) commonly become nature reserves instead (Statistics Sweden 2023). Nature conservation agreements, however, have no practiced upper size limit and can be established, within the same legal format, on larger areas such as the Sveaskog State forest company ecoparks (Bergman & Gustafsson 2020) with Tjades-Nimtek ecopark in the western parts of the north boreal region being the largest with 22.300ha (Sveaskog, 2021).

In Sweden, identification and registration of a woodland key habitat (WKH), defined as an area with high forest conservation value (Timonen et al. 2010), is a basis for NCA and BPA establishments (Statistics Sweden 2018). The inventory and registration of WKHs in Sweden has been subject to a heated debate (Jakobsson et al. 2021), resulting in suspension in 2019 as the Swedish government withdrew the funding from the Swedish Forest Agency (Angelstam et al. 2020). Hence, reliable data dates some years back, but clearly indicates that high conservation forests occur at a substantial share on NIPF-ownership. Up to 2015, the Swedish Forest Agency register included around 100,000 WKHs covering 470,000 ha on national scale, whereof 61,200 WKHs covering 170,000 ha outside the large forest company ownership (Wester & Engström 2016). The voluntary forest certification (Lehtonen et al. 2021) requires that landowners set-aside forest with high conservation value, and while many NIPF-owners are not certified, a large share of all harvesting is done by actors that operate under certification schemes, i.e. by private forest owners associations and State and private forest industry companies. The turbulence around and suspension of the WKH-inventory may have contributed to the drop in the establishment of NCAs and BPAs since 2006. This is unfortunate as it risks loss of forests with high conservation value through thinning operations, since only final clear-cut harvesting is regulated and restricted according to the Swedish Forestry Act (Jonsson et al. 2019).

Commercial forestry in Sweden is largely restricted to Scots pine and Norway spruce (Svensson et al. 2023a). To some extent, birch, aspen and the introduced non-native Lodgepole pine, and for south Sweden also oak and beech, contributes to the forest production scheme of tree species. Spruce, pine and Lodgepole pine in single-species and mixed stands cover 82% of the forestlands in use for commercial forestry in Sweden, whereof spruce contributes 27% (SLU 2023). Interestingly, removal of spruce dominates totally in restoration oriented towards removing dis-favored tree species, both in young, plantation forests and in older forests with merchantable trees. Favored tree species, on the other hand, include in total 24 different tree and tall woody shrub species, although with a few species clearly dominating. Evidently, a restoration ambition is clearly to diversify species composition on stand and landscape level. It should be noted that several of the 24 species are species with no or low commercial wood value, but that contribute other qualities and functions to forest ecosystems.

Our results show that representative forest types are protected, with deciduous hardwood forests dominating in south Sweden, and spruce, pine and mixed coniferous forests in the north. Compared with forest statistics (SLU, 2023), however, NCA and BPA cover a larger proportion of deciduous (15.7% vs. 6.5%), deciduous hardwood forests (9.8% vs. 1.1%) and mixed forests (12.6% vs. 6.9%), relative to coniferous forests (62% vs. 82%), indicating a protections strategy oriented towards less abundant forest types on national scale. The SLU (2023) proportions presented here, i.e. the latter figure within the brackets, concern productive forest of interest for commercial forestry and combines pine, spruce, Lodgepole pine (*Pinus contorta*) and mixed coniferous forests.

Our results also shows that active restoration, i.e. the combined area of any type of restoration management, exceeds passive restoration, i.e. area set aside for natural, undisturbed development, in the NCAs and BPAs; 57% vs. 43%. Our finding here clearly exceeds previous estimates; Wester and Engström (2013) reported that a share equal to 13% of the WKH-area was in need for active restoration management. Moreover, our findings shed needed light on the fact that management restoration interference is needed to put also high conservation value forests in a better ecological state than at present, across and extensive biogeographical gradient of northern European forest landscapes.

Collectively, the restoration types analyzed well represents the range of restoration options known and implemented for protection forest biodiversity (Bernes et al. 205). Setting aside areas are the dominant form of restoration in the south and north boreal ecoregions, reflecting larger opportunities to build on existing fragments of natural and near natural forests with high current conservation value (e.g., Bubnicki, et al. 2024). Among forest-stand restoration management types, removal of tree species and thinning from above are the most abundant types across all ecoregions, whereas prescribed burning and creation of dead wood are most abundant in the boreonemoral and the south and north boreal ecoregions. An overall reflection is that the applied restoration management is concentrated to a fairly low number of different types. Given the natural variability in forest configuration and dynamics across the northern European ecoregions, we foresee that there is potential to diversify restoration further. Such a diversification should consider aspects such as different duration and degree of land-use, different land ownership situations, multiple-use of forests, and adaptation to climate change as well as to emerging new markets and value chains (cf. Gann et al. 2019).

In Sweden, land use on NIPF-ownership is similar to large company ownership and strongly dominated by wood biomass yield based on systematic rotation forestry (e.g., Jonsson, et al. 2019). In parallel, other important values and land use interests exists, such as indigenous Sami people reindeer husbandry in northern Sweden (Pape & Löffler 2012), recreation, tourism and other interests associated with the public right of access (Saito et al. 2023; Haukeland et al. 2023), and hunting (Neumann et al. 2022). With other land uses and market chains expanding on forestland, e.g. wind power establishments (Svensson et al. 2023b), and carbon- and biodiversity credits (Iwarsson Wide 2022; Le et al. 2024.), it can be assumed that forest governance and management directions will diversify (Knoke et al. 2017; Felton et al. 2020). As a result, land use will develop from single use to multiple use more generally. Hence, restoration will be needed as a key component of the forestry model to put forest habitats in a state that secure biodiversity and functional ecological interactions, but also allow provisioning of multiple goods and services (Ferraro et al. 2013; Vallecillo et al. 2018; Svensson et al. 2023a).

It is positive that the agreements include clear management objectives and guidelines on how often actions should be performed. The time interval between activities is generally short, i.e. commonly up to five or ten years regardless of type, with exception of prescribed burning that re-occurs at all intervals up to 50 years. Although recognizing that restoration generally is expected to re-occur frequently, and thus need to be as frequently planned, executed and financed, we, however, question the time interval relevance given internal dynamics and natural disturbances in forest ecosystems (cf. Kuuluvainen & Aakala, 2011; Berglund & Kuuluvainen, 2021). Thinning forest stands, creating small gaps and promoting tree regeneration of favored species represent activities that may need to be repeated frequently. However, for other activities, the recurrence interval seems less valid. For example, prescribed burning or creating deadwood can be done once or in long time intervals, i.e. longer than 50 years, given longevity of trees and resilient ecological processes in boreal forest ecosystems (e.g., Esseen et al. 1997). Furthermore, prescribed burning is hardly relevant in these small-size forests with an interval of a few years or even a few decades (cf. Nicklasson & Granström, 2000). We find that time interval between restoration management activities will need further exploration. A major question though, relates to whether these activities are actually executed and evaluated concerning their nature conservation benefits and their importance for securing functional ecological processes and interactions. No such information is currently available. Ideally, an adaptive management approach with defined management actions following regular monitoring, should be developed.

The potential of NCA and BPA remains an option for economic compensation for lost forest harvesting opportunities due to certification rules, but naturally also for attracting interest among landowners to voluntary, and with a certain amount of economic compensation, set aside forest for other purposes that wood biomass yield. Emerging economic diversification (Knoke et al. 2017) including alternative value chains such as carbon- and biodiversity credits (Iwarsson Wide 2022), offer similar types of agreements as NCAs, as being voluntary and often agreed for certain restricted time periods, but with funding from external actors besides from the State. As the need for forest and forest landscape restoration is high in Sweden and elsewhere where the industrial footprint is substantial, we argue that both carbon- and biodiversity credits on an open market may complement State investment in the form of NCAs and BPAs to promote a higher nature conservation and environmental ambition, in particular among NIPF-owners. It is clear that value chains and business models for forest owners need to develop in parallel with expanded forest protection and restoration. It is critical, however, that these emerging financing instruments provide additionally and consider leakage effects to actually contribute to forest conservations (Wunder et al., in prep.).

Since NCAs can be established based on other aspects than high nature conservation values (Statistics Sweden 2018), this indicates a potential for developing such formal agreements as a business model for promoting a forest management system that favors, for example, recreational values, conservation of cultural heritage, good grazing conditions for reindeer, i.e. promoting forest multiple use aspects. Up to 2023, however, only 15 NCAs covering in total 87 ha has been established for other values than nature conservation (Swedish Forest Agency 2024). Here, we argue that there is an evident potential to be further explored. The NCA-designation, and potentially also restoration components of the more strict BPA biodiversity conservation construction as a legal instrument, provide opportunities to support diverse forest values through restoration, providing alternative targets for landowners.

Furthermore, the dialogue between the State and the landowner to reach a voluntary agreement, showcase a concrete way to engage stakeholders in participatory governance, which is a principle of forest landscape restoration (Besseau et al. 2018). We argue that NCAs, conceptually, exemplifies a formal instrument that merits further exploration, in particular if restoration management imply an opportunity for the landowner to harvest and sell trees, simultaneously as supporting other values that society demands (cf. Knoke et al. 2017). There is a vast potential for restoration and conservation in NCAs and BPAs in Sweden, which is currently underutilized and has further value towards development of any biodiversity credit system under development. In a landscape context, spatial planning based on selection of critical forest types for protection and restoration, and of landowners that are motivated, can potentially form a network of small-size protection areas that support ecological connectivity. With emphasis on restoration, this approach would accentuate the key question of where to restore to achieve expected ecological response (Wang et al. in prep.).

To meet nature conservation, biodiversity and environmental goals, specifically, restoration will be needed for improving the forest ecosystem status, including transformation of forest configuration to secure natural biotic and abiotic processes in functional forest ecosystems on stand and landscape scale. In countries where systematic, industrial forestry have left a too strong anthropogenic footprint on the forest landscapes, such as in Sweden, restoration will need to be extensive, well informed and flexible based on, e.g., the landowner situation and the landowners capacity, interest and choice. With non-industrial private forest owners being dominant, good examples on incentives for restoration are of high value. Here, restoration approaches in NCAs and BPAs provide a rich source of experiences. In counties and regions with fragmented private landownership, sustainable conservation planning and restoration of representative, functional and well-managed forests for biodiversity protection, include small-size forest areas (e.g., Fahrig 2020) with or without mitigation activities to reduce negative consequences on biodiversity and functional connectivity (Brennan et al. 2022). The gradient explored in this study, from the nemoral ecoregion in the southernmost to the north boreal and northwest subalpine in northernmost Sweden, clearly shows the importance of integrating small protected areas in a national conservation scheme. In south Sweden, NCAs and BPAs contribute substantial protected areas, but not in north Sweden. Since large protected areas are, basically, restricted to the Scandinavian Mountain Green Belt forest foothills landscape (Svensson et al. 2020), the relative low occurrence of NCAs and BPAs on remaining natural and near natural forests in the heavily transformed coastal and inland forest landscapes of north Sweden (Bubnicki et al. 2024), is unfortunate. Especially in the light of the EU nature restoration law, models and instruments are needed to provide private forest owners with incentives that include protection of and initiate restoration in small-size forests with high conservation and multiple use values and high values. Here the NCAs and BPAs represent an example of an instrument that can be further developed and implemented.

## Acknowledgement

This study is part of the SUPERB: Upscaling forest restoration EU Horizon 2020 Research and Innovation Programme (SUPERB: Upscaling Forest Restoration - SUPERB (forest-restoration.eu); Grant Agreement number 101036849). We acknowledge the assistance by Göte Eriksson, Swedish Forest Agency, for compiling and providing the database with nature conservation agreements and biotope protection areas in Sweden.

## Author contributions

JS, ALP, BGJ and NJS share first authorship. JS conceived the idea, scope and context, and coordinated the analyses, writing and design. ALP compiled data and performed analyses, designed and developed the illustrations. ALP, BGJ and NJS participated in scope development, interpretation of results, and writing.

