## Supplementary material for "The contribution of small protected areas to the big picture of forest conservation and restoration in Sweden"

**S1**. Restoration management types and categories

**Table S1**. Restoration management types in the database includes in total 30 different categories, whereof 13 combined categories were constructed.

| Restoration management categories | Restoration management types included |
| --- | --- |
| Forest floor and hydrology | - Measure in field- and bottom layer - Top soil disturbance - Restore hydrology - Restore watercourses |
| Shrub layer | - Cleaning in shrub layer - Favoring shrub layer |
| Tree regeneration | - Tree-specific regeneration Management - Cleaning in tree regeneration |
| Tree age heterogeneity | - Increase tree age heterogeneity by selective harvesting |
| Removal of tree species | - Removal of tree species by selective harvesting |
| Thinning from above | - Favor selected tree species and tree groups in canopy layer - Open up canopy layer to favor selected tree species and groups |
| Gap cutting | - Gap cutting |
| Dead wood | - Increase amount of dead wood |
| Prescribed burning | - Prescribed burning |
| Forest edge | - Maintain or restore forest edge - Maintain or restore forest edge towards water |
| Cultural and recreational values | - Livestock grazing in forest habitats with disturbance-favored values - Pollarding of trees - Management of cultural values - Mowing - Recreational values |
| Other | - Other, not specified - Species-specific management activities - Increase overall heterogeneity in the area - Other single-tree specific management - Other, miscellaneous measures and management - Other, key element/attribute measures - Management of specifically valuable trees |
| Set aside | - Set aside for natural, free development |

**S2**. Area of Nature Conservation Agreements and Biotope Protection Areas

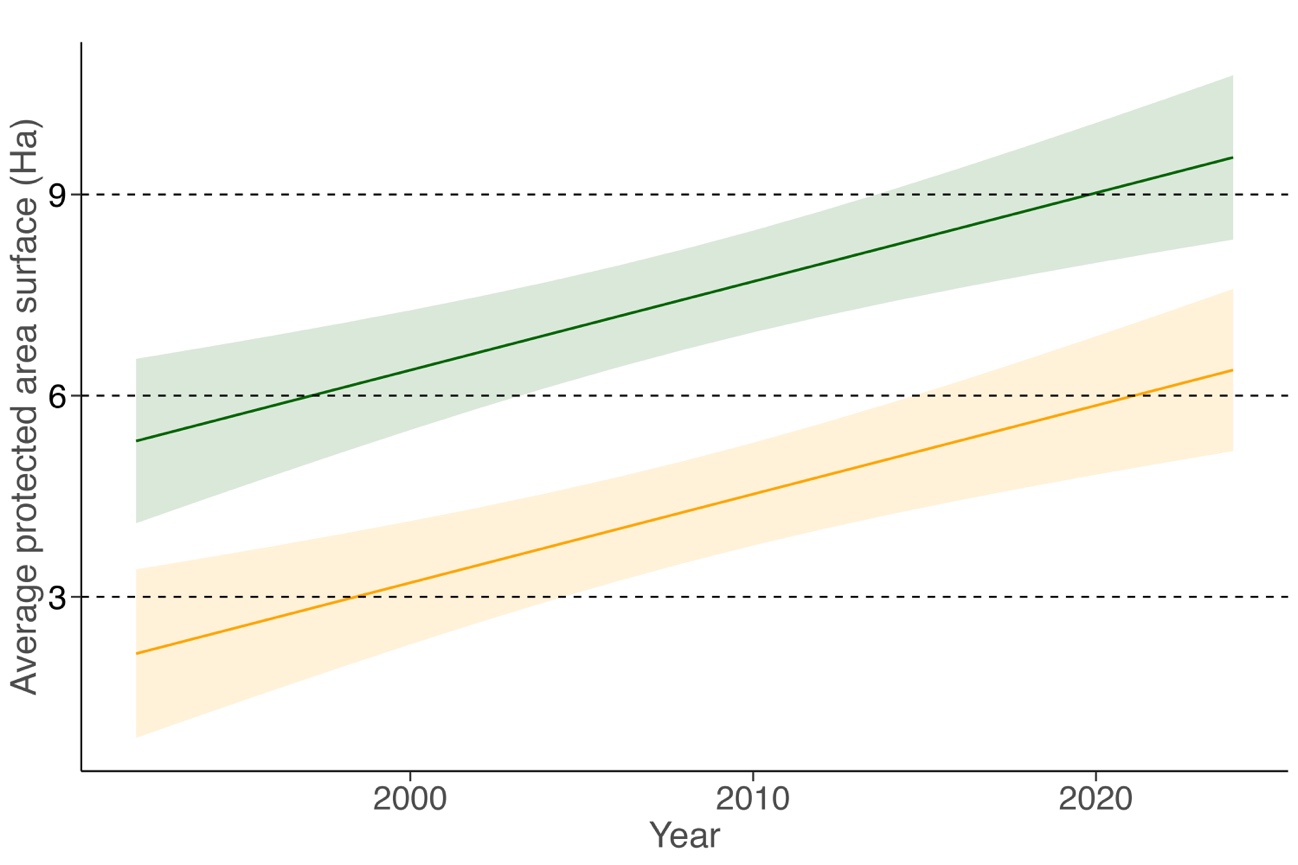

**Figure S2**. Predicted increase in total area (ha) of all Nature Conservation Agreements (from the first established October 1, 1993) in green, and all Biotope Protection Areas (from the first established February 28, 1994), in yellow, up to 2023 (June 6). The line shows predicted increase in protected area surfaces with the shaded areas indicating the 95% Confidence interval. The average size is increasing significantly over time; NCA estimate = 0.13, t = 4.36, p <0.01, and BPA estimate = 1.59, t = 5.94, p <0.01.

**S3**. Forest type area in Nature Conservation Agreements and Biotope Protection Areas

**Table S3**. Area (ha) of the six different forest types in all Nature Conservation Agreements (from the first established October 1, 1993) and all Biotope Protection Areas (from the first established February 28, 1994) up to 2023 (June 6), combined*, in database provided by the Swedish Forest Agency.

| Ecoregion | Forest type | Not on wetland (ha) | On wetland (ha) | Total area (ha) |
| --- | --- | --- | --- | --- |
| North boreal | Pine forest | 3,704 | 202 | 3,906 |
|  | Spruce forest | 4,410 | 124 | 4,533 |
|  | Mixed coniferous forest | 1,143 | 23 | 1,165 |
|  | Mixed forest | 1,818 | 174 | 1,991 |
|  | Deciduous forest | 539 | 118 | 657 |
|  | Deciduous hardwood forest* | - | - | - |
| South boreal | Pine forest | 7,843 | 878 | 8,721 |
|  | Spruce forest | 9,061 | 267 | 9,328 |
|  | Mixed coniferous forest | 3,811 | 155 | 3,966 |
|  | Mixed forest | 2,593 | 488 | 3,081 |
|  | Deciduous forest | 3,393 | 467 | 3,860 |
|  | Deciduous hardwood forest* | 394 | 0 | 394 |
| Boreonemoral | Pine forest | 6,096 | 817 | 6,913 |
|  | Spruce forest | 4,552 | 130 | 4,682 |
|  | Mixed coniferous forest | 2,728 | 97 | 2,824 |
|  | Mixed forest | 3,400 | 432 | 3,832 |
|  | Deciduous forest | 5,243 | 1,283 | 6,525 |
|  | Deciduous hardwood forest* | 4,180 | 5 | 4,185 |
| Nemoral | Pine forest | 386 | 38 | 424 |
|  | Spruce forest | 277 | 10 | 287 |
|  | Mixed coniferous forest | 102 | 3 | 105 |
|  | Mixed forest | 573 | 44 | 618 |
|  | Deciduous forest | 723 | 122 | 846 |
|  | Deciduous hardwood forest* | 2,867 | 1 | 2,868 |

* Combined NLC-categories “hardwood deciduous forest” and “hardwood deciduous forest with trivial deciduous forest” (Swedish Environmental Protection Agency 2018). Dash (-) shows no occurrence, zero (0) shows occurrence below 0.5ha.

**S4**. Area of restoration management types

**Table S4**. Area (ha) of the 13 different restoration management types in all Nature Conservation Agreements (from the first established October 1, 1993) and all Biotope Protection Areas (from the first established February 28, 1994) up to 2023 (June 6), combined, in database provided by the Swedish Forest Agency.

| Ecoregion | Restoration Management type | Total area (ha) |
| --- | --- | --- |
| North boreal | Forest floor and hydrology | 37 |
|  | Shrub layer | 1 |
|  | Tree regeneration | 292 |
|  | Tree age heterogeneity | 4 |
|  | Removal of tree species | 311 |
|  | Thinning from above | 521 |
|  | Gap cutting | 84 |
|  | Dead wood | 379 |
|  | Prescribed burning | 1,391 |
|  | Forest edge | 13 |
|  | Cultural and recreational values | 43 |
|  | Other | 548 |
|  | Set aside | 8,011 |
| South boreal | Forest floor and hydrology | 173 |
|  | Shrub layer | 60 |
|  | Tree regeneration | 470 |
|  | Tree age heterogeneity | 104 |
|  | Removal of tree species | 2,715 |
|  | Thinning from above | 1,669 |
|  | Gap cutting | 161 |
|  | Dead wood | 1,135 |
|  | Prescribed burning | 1,916 |
|  | Forest edge | 33 |
|  | Cultural and recreational values | 448 |
|  | Other | 1,474 |
|  | Set aside | 13,830 |
| Boreonemoral | Forest floor and hydrology | 166 |
|  | Shrub layer | 882 |
|  | Tree regeneration | 1,392 |
|  | Tree age heterogeneity | 549 |
|  | Removal of tree species | 4,956 |
|  | Thinning from above | 4,945 |
|  | Gap cutting | 768 |
|  | Dead wood | 1,683 |
|  | Prescribed burning | 298 |
|  | Forest edge | 108 |
|  | Cultural and recreational values | 3,649 |
|  | Other | 1,781 |
|  | Set aside | 7,734 |
| Nemoral | Forest floor and hydrology | 10 |
|  | Shrub layer | 157 |
|  | Tree regeneration | 569 |
|  | Tree age heterogeneity | 41 |
|  | Removal of tree species | 1,439 |
|  | Thinning from above | 1,393 |
|  | Gap cutting | 60 |
|  | Dead wood | 187 |
|  | Prescribed burning | 6 |
|  | Forest edge | 39 |
|  | Cultural and recreational values | 477 |
|  | Other | 1,575 |
|  | Set aside | 1,010 |

**S5.** Favored tree species

**Table S5**. Favored tree species in subtype Tree specific regeneration management within restoration type 3 (Tree regeneration) and in subtype Favor selected tree species and tree groups in canopy layer within restoration type 6 (Thinning from above), area (ha). Category other include areas within the sub types where specific species are not defined

| Species | Latin name | Area (ha) Canopy | Area (ha) Understory |
| --- | --- | --- | --- |
| Alder | Alnus glutinous, incana | 157 | - |
| Ash | Fraxinus excelsior | 228 | 24 |
| Aspen | Populus tremula | 1,606 | 384 |
| Beech | Fagus sylvatica | 99 | 51 |
| Bird cherry | Prunus padus | 3 | 1 |
| Birch | Betula pendula, pubescens | 973 | 169 |
| Buckthorn | Frangula alnus | - | 4 |
| Conifer trees |  | 1 | 19 |
| Deciduous trees |  | 170 | 43 |
| Elm | Ulmus glabra, etc. ssp | 118 | 4 |
| Hazel | Coryllus avellana | 200 | 20 |
| Hornbeam | Carpinus betulus | 4 | 7 |
| Juniper | Juniperus communis | 43 | 3 |
| Lime | Tilia cordata, etc. ssp | 282 | 16 |
| Maple | Acer platanoides, etc. ssp | 199 | 3 |
| Oak | Quercus robur, petraea | 2,424 | 97 |
| Pine | Pinus sylvestris | 1,567 | 238 |
| Rowan | Sorbus aucuparia | 126 | 161 |
| Spruce | Pive abies | 82 | 7 |
| Wild apple | Malus spp | 57 | - |
| White beam | Sorbus intermedia | 21 | 3 |
| Wild cherry | Prunus avium | 53 | - |
| Willow | Salix spp | 461 | 177 |
| Yew | Taxus baccata | 1 | - |
| Subtotal |  | 8,874 | 1,428 |
| Other |  | 1,989 | 282 |
| Total |  | 10,863 | 1,710 |

**S6.** Dis-favored tree species

**Table S6**. Dis-favored species, extracted from restoration type Removal of tree species, area (ha). Category other include areas within the sub types where specific species are not defined, dbh = diameter at 1.3m

| Dis-favored species | Area (ha) |
| --- | --- |
| Harvesting of non-native tree species >15 cm dbh | 69 |
| Harvesting of spruce >15 cm dbh | 2,552 |
| Harvesting of deciduous tree species >15 cm dbh | 61 |
| Harvesting of pine >15 cm dbd | 7 |
| Cleaning of non-native tree species | 41 |
| Cleaning of spruce | 4,744 |
| Cleaning of deciduous trees | 186 |
| Cleaning of pine | 12 |
| Subtotal area | 7,671 |
| Other | 281 |
| Total area | 7,952 |
